# Spheroplast-mediated carbapenem tolerance in Gram-negative pathogens

**DOI:** 10.1101/578559

**Authors:** Trevor Cross, Brett Ransegnola, Jung-Ho Shin, Anna Weaver, Kathy Fauntleroy, Michael VanNieuwenhze, Lars F. Westblade, Tobias Dörr

## Abstract

Antibiotic tolerance, the ability to temporarily sustain viability in the presence of bactericidal antibiotics, constitutes an understudied, yet likely widespread cause of antibiotic treatment failure. We have previously shown that the Gram-negative pathogen *Vibrio cholerae* is able to tolerate exposure to the typically bactericidal β-lactam antibiotics by assuming a spherical morphotype devoid of detectable cell wall material. However, it is unclear how widespread tolerance is. Here, we have tested a panel of clinically significant Gram-negative pathogens for their response to the potent, broad-spectrum carbapenem antibiotic meropenem. We show that clinical isolates of *Enterobacter cloacae, Klebsiella pneumoniae*, and *Klebsiella aerogenes*, but not *Escherichia coli*, exhibit moderate to high levels of tolerance to meropenem, both in laboratory growth medium and in human serum. Importantly, tolerance was mediated by cell wall-deficient spheroplasts, which readily recovered to wild-type morphology and exponential growth upon removal of antibiotic. Our results suggest that carbapenem tolerance is prevalent in clinically significant bacterial species, and we suggest that this could contribute to treatment failure associated with these organisms.

## Introduction

Antibiotics are often differentiated by their ability to either inhibit growth (bacteriostatic) or to kill bacteria (bactericidal). The exact differentiation of antibiotics into these broad categories likely depends on the species and the specific growth environment in which antibiotic susceptibility is tested (1, 2). To optimize therapy, it is essential to gain a comprehensive understanding of the variable factors that modulate bacterial susceptibility to antibiotics. For example, the β-lactams (penicillins, cephalosporins, cephamycins, carbapenems, and the monobactam aztreonam), which are among the most powerful agents in our antibiotic armamentarium, prevent and/or corrupt proper cell wall (peptidoglycan) assembly (3, 4). Consequently, these agents typically induce cell death and lysis in susceptible bacteria at least during rapid growth *in vitro* (3, 4). However, *in vivo* β-lactams often fail to eradicate an infection caused by susceptible (*i.e*., non-resistant) organisms (5–7). This paradox can, in part, be explained by the presence of dormant persister cells, a small subpopulation that resists killing by antibiotics that require cellular activity for their lethal action (8–10). However, specimens obtained from patients treated with β-lactam antibiotics have been reported to contain spheroplasts, bacterial cells that lack a cell wall (11)(3), and clinical isolates are often highly tolerant to β-lactam antibiotics at frequencies that cannot solely be explained by invoking rare persister cells (4,5)(12). Spheroplast formation suggests that in these bacteria the antibiotic is effective in inhibiting cell wall synthesis, demonstrating that some bacteria survive antibiotic exposure in forms that are neither dormant nor resistant. We and others have previously shown that two important Gram-negative pathogens, *Vibrio cholerae* and *Pseudomonas aeruginosa*, form viable, non-dividing spheroplasts when exposed to inhibitors of cell wall synthesis (6,7)(13). Spheroplasts readily revert to rod-shape and exponential growth, suggesting these cells might promote re-infection upon discontinuation of antibiotic therapy. Successful recovery of *V. cholerae* spheroplasts requires the cell wall stress sensing two-component system VxrAB (also known as WigKR (14, 15)), cell wall synthesis functions and the general envelope stress-sensing alternative sigma factor RpoE (16).

Spheroplast formation is reminiscent of so-called Gram-positive “L-forms”: irregularly dividing, cell wall-less cells surrounded only by their cytoplasmic membranes (9). However, in striking contrast to L-forms, Gram-negative spheroplasts do not divide in the presence of antibiotic (13, 17). Division through an L-form-like mechanism is likely prevented by the presence of their strong outer membrane (OM) which exhibits almost cell wall-like mechanical properties (18). Indeed, dividing L-forms of the Gram-negative model organism *Escherichia coli* can be generated by inhibiting cell wall synthesis in osmostabilized growth medium, which causes the cytoplasm to “escape” its OM shell (19).

While L-form formation has been implicated as a mechanism of antibiotic resistance in Gram-positive bacteria (11), it is unclear whether spheroplast formation represents a general strategy elicited by Gram-negative bacteria to tolerate cell wall synthesis inhibitors such as the β-lactams. Here, we have tested a collection of well-characterized American Type Culture Collection (ATCC) and clinical Gram-negative isolates (Table 1) for their ability to tolerate exposure to the carbapenem antibiotic meropenem. We find that, with the notable exception of *E. coli*, all isolates formed cell wall-deficient spheroplasts upon exposure to meropenem, and these spheroplasts were able to fully recover to rod-shape and exponential growth upon removal of meropenem, both in a laboratory medium and in human serum. Our data suggest that spheroplast-mediated carbapenem tolerance is prevalent in clinically significant Gram-negative pathogens, but rare or absent in the *E. coli* isolates tested herein. Our results suggest that measures of antibiotic susceptibility and ultimately treatment outcome could consider more nuanced responses in diverse, clinically-relevant Gram-negative pathogens.

**Table 1.**
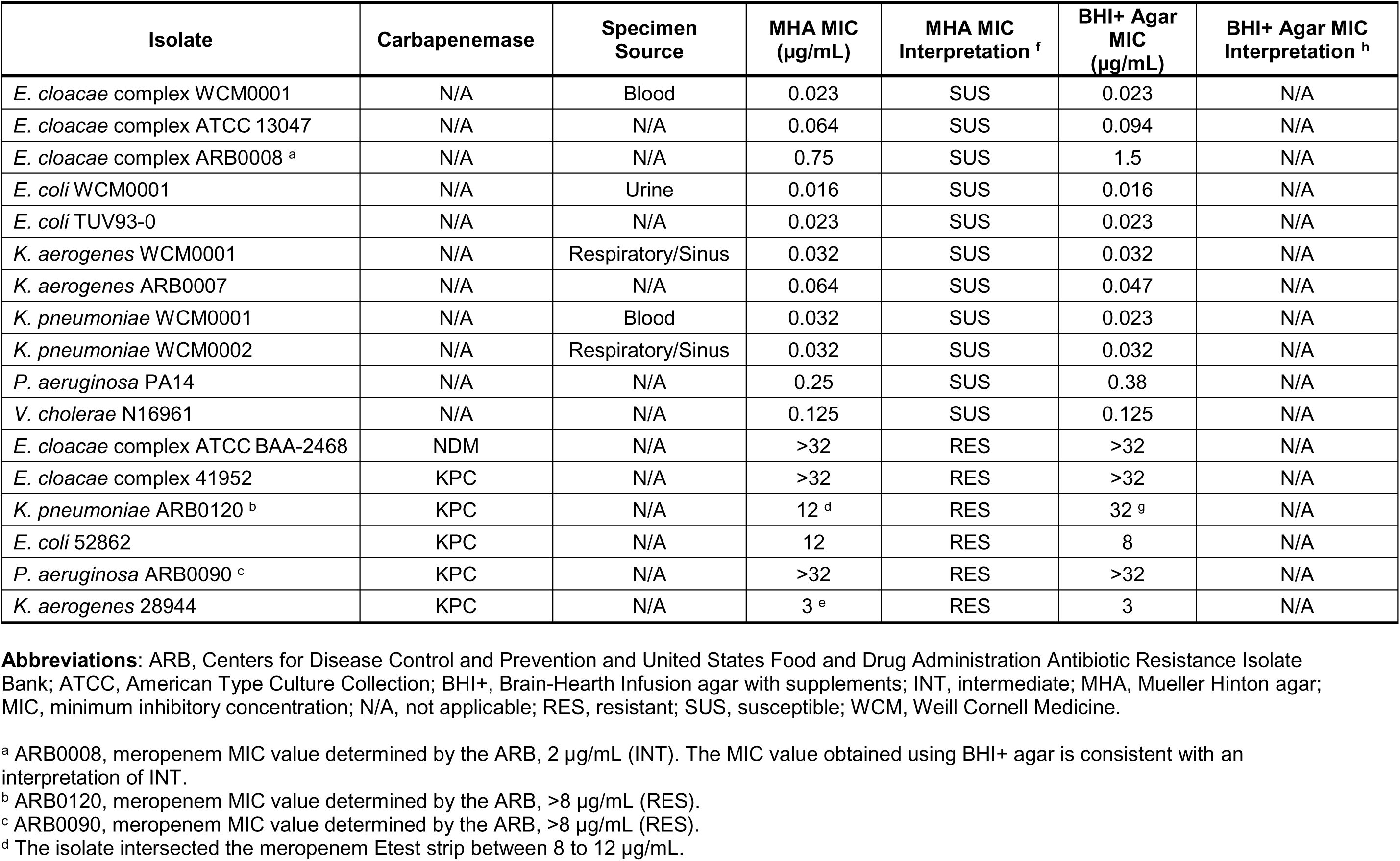
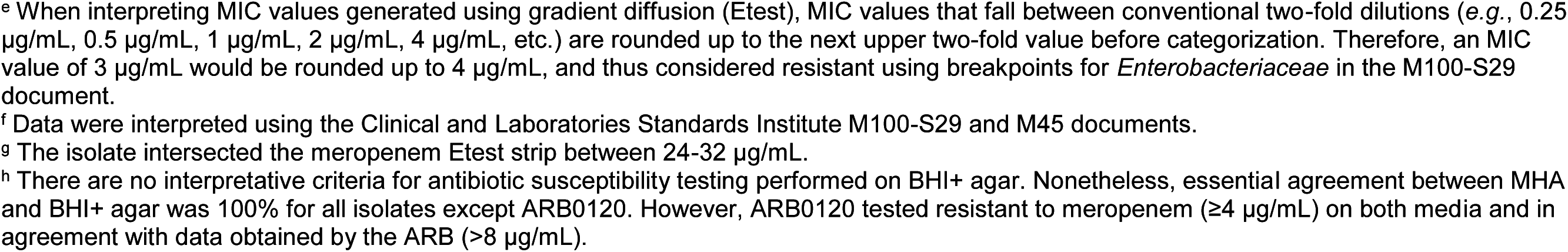
Bacterial isolates evaluated in this study.

## Results

### Tolerance to Meropenem Varies Across Gram-Negative Clinical Isolates

Spheroplast-mediated β-lactam tolerance might be an underappreciated menace in the clinical setting. To test how widespread the ability to tolerate cell wall acting antibiotics is in clinical isolates, we assayed a panel of clinical isolates representative of significant Gram-negative pathogens of the family *Enterobacteriaceae*: *E. coli*, including Enterohemorrhagic *E. coli* (EHEC), *Enterobacter cloacae*, *Klebsiella aerogenes* (formerly *Enterobacter aerogenes*), and *Klebsiella pneumoniae*. We also tested organisms known to form spheroplasts under some conditions: *V. cholerae* and *P. aeruginosa*. As a representative β-lactam, we used the carbapenem meropenem. We chose meropenem due to its importance as a potent, broad-spectrum agent (20, 21), and also because in clinical practice, especially in the setting of multi-drug resistance, it is often used against members of our isolate panel (20, 22).

We conducted time-dependent killing experiments measuring both colony forming units (cfu/ml) and optical density (OD_600_). Killing experiments for all isolates were conducted in supplemented Brain Heart Infusion (BHI+) broth under high inoculum/slow growth conditions (see Methods for details) to emulate the likely slow growth behavior during an infection (23). We chose a meropenem concentration (10 µg/mL), which is above the meropenem resistant breakpoint for *Enterobacteriaceae* [≥4 µg/mL], *P. aeruginosa* [≥8 µg/mL], and *V. cholerae* [≥4 µg/mL] (24, 25) and between 6.7 × and 625 × higher than the minimum inhibitory concentration (MIC) for each susceptible/non-resistant, non-carbapenemase producing isolate (Table 1). Crucially, antibiotic susceptibility testing (AST) revealed that MIC values did not differ significantly between media Mueller Hinton agar [MHA] (recommended for AST by the Clinical and Laboratories Standards Institute (CLSI) (24, 25)) and BHI+ agar; the essential agreement (26) between MIC values on both media was 100% for all but one isolate (ARB0120, although the isolate exhibited MIC values greater than the meropenem resistant breakpoint on both MHA and BHI+ agar) (Table 1), suggesting that AST performed with BHI+ is comparable with standardized methods. For comparison to the susceptible/non-resistant strains, we included a panel of conspecific clinical isolates that are carbapenem-resistant due to their possession of the carbapenemase KPC (***K**lebsiella **p**neumoniae* **c**arbapenemase).

Among the susceptible/non-resistant, non-carbapenemase producing isolates, killing and optical density dynamics varied widely between species and even isolates within the same species (*e.g*., *E. cloacae* WCM0001 versus *E. cloacae* ARB0008) (Fig. 1). Interestingly, both in lysis behavior and survival, *E. coli* was considerably less tolerant than all other tested organisms (Fig. 1). While killing efficiency after 6 hours of meropenem exposure generally ranged from ~ 5 to10-fold killing (*V. cholerae* N16961*, P. aeruginosa* PA14*, E. cloacae* WCM0001, *E. cloacae* ATCC 13047) to ~5,000-fold killing (*K. aerogenes* WCM0001*, E. cloacae* ARB0008, *K. pneumoniae* WCM0001 and WCM0002), both *E. coli* isolates tested were almost completely eradicated by meropenem (~10^8^-fold killing) (Fig. 1, **S1 - S2**). In contrast, almost all isolates grew well in the absence of meropenem, except for both isolates of *P. aeruginosa*, which exhibited slower growth in BHI+ compared to the other isolates (**Fig. S3**). Interestingly, *E. cloacae* ARB0008 had a higher meropenem MIC than the other *E. cloacae* isolates, but was among the isolates with the highest degree of killing. This observation suggests that susceptibility (*i.e*., differences in MIC values) might not necessarily correlate with tolerance, *i.e*., the degree of killing.

**Figure 1.**
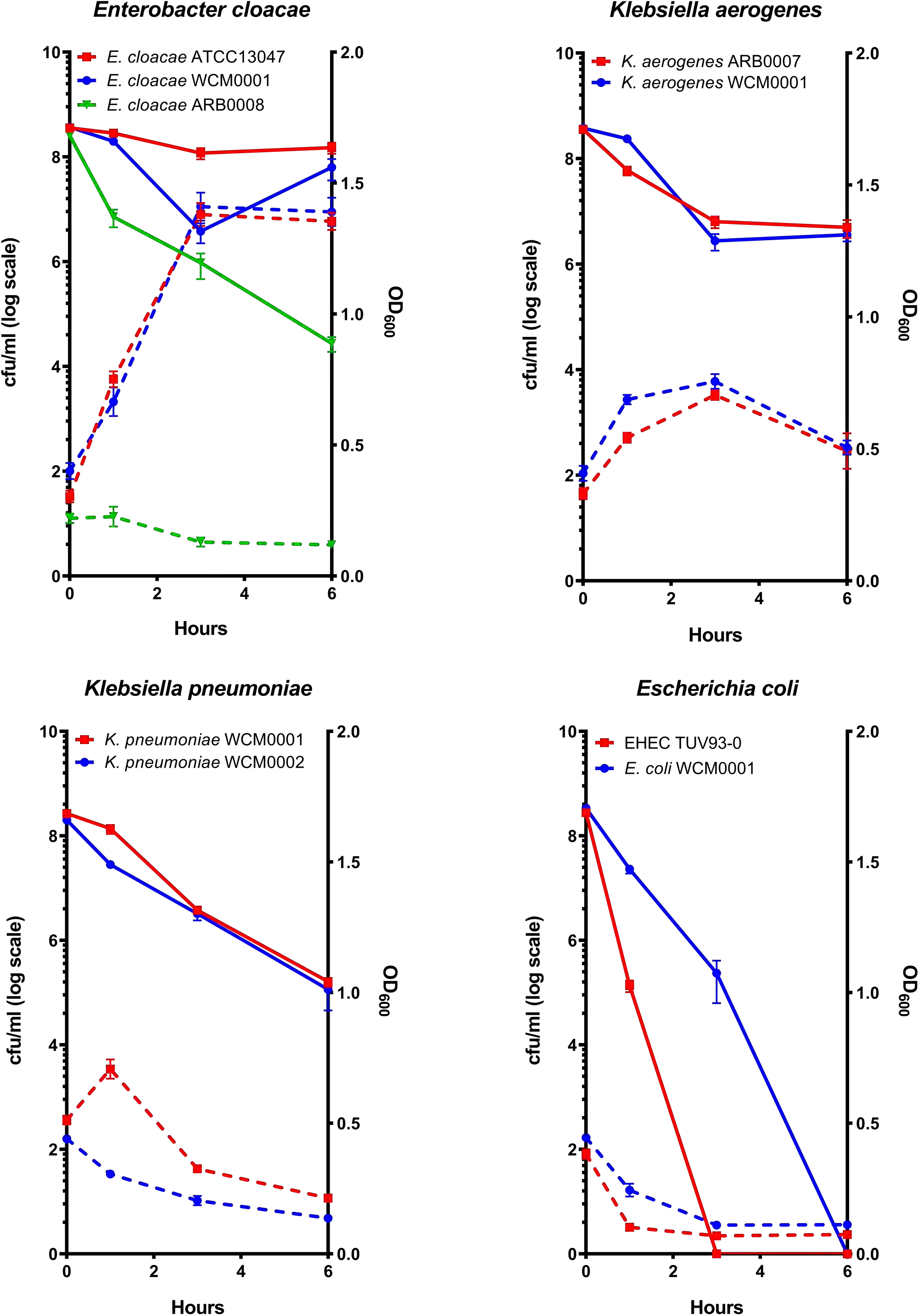
Clinical Gram-negative pathogens exhibit variable degrees of killing after exposure to meropenem. Overnight cultures of the indicated isolates were subcultured 1:10 (final volume 5 mL) into pre-warmed BHI+ liquid medium supplemented with 10µg/mL meropenem. Optical density (OD_600_, dotted lines) and viable cell counts (cfu/mL, solid lines) were measured at the indicated time points. Error bars represent standard error of the mean of at least six biological replicates.

Similar to *V. cholerae* and *P. aeruginosa* (**Fig. S1 – S2** and (13)), survival of meropenem-treated cells often coincided with a substantial increase in OD_600_ (while cfu/mL stayed the same or decreased)(Fig. 1), demonstrating that surviving cells are not dormant, but as a population continue to increase in mass. This was not observed in either isolate of *E. coli*, where a rapid decrease in OD_600_ indicated lysis during meropenem exposure. Thus, meropenem tolerance (and potentially β-lactam tolerance in general) is appreciable in clinical isolates of some *Enterobacteriaceae*, but not observed (at least under conditions tested herein) in *E. coli*. In comparison to the susceptible isolates, and as expected, all KPC positive isolates increased in both OD_600_ and cfu/ml during meropenem exposure (**Fig. S4**).

### Meropenem tolerant survivors are cell wall-deficient spheroplasts

In *V. cholerae* and *P. aeruginosa*, β-lactam tolerant cells are cell wall-less, metabolically active spheroplasts. In principle, the moderate to high level tolerance we observed in our experiments could also be a consequence of unusually high levels of dormant persister cells in these clinical isolates due to a prolonged lag phase (27) emerging from stationary phase. To distinguish between these two possibilities, we withdrew samples at various time points following exposure to meropenem and imaged them. Dormant persister cells remain rod-shaped in the presence of cell wall acting antibiotics, since these cells prevent antibiotic damage completely through their lack of growth (28, 29). Visual examination revealed that, comparable to previous observations in *V. cholerae* and *P. aeruginosa* (6,7), the tolerant populations of almost all isolates consisted exclusively of spherical cells (Fig. 2 and **S1-S2, S5 – S8**). The notable exception were the two *E. coli* isolates; while some spherical cells could be observed after short exposure periods in both tested isolates, after 6 hours of antibiotic exposure only cell debris was observed (Fig. 2 and **Fig. S9**). In contrast to those cultures treated with meropenem, untreated bacteria retained rod-shape in BHI+ (**Fig. S10**); similar to the conspecific, but KPC positive, isolates with or without meropenem treatment (**Fig. S11**).

**Figure 2.**
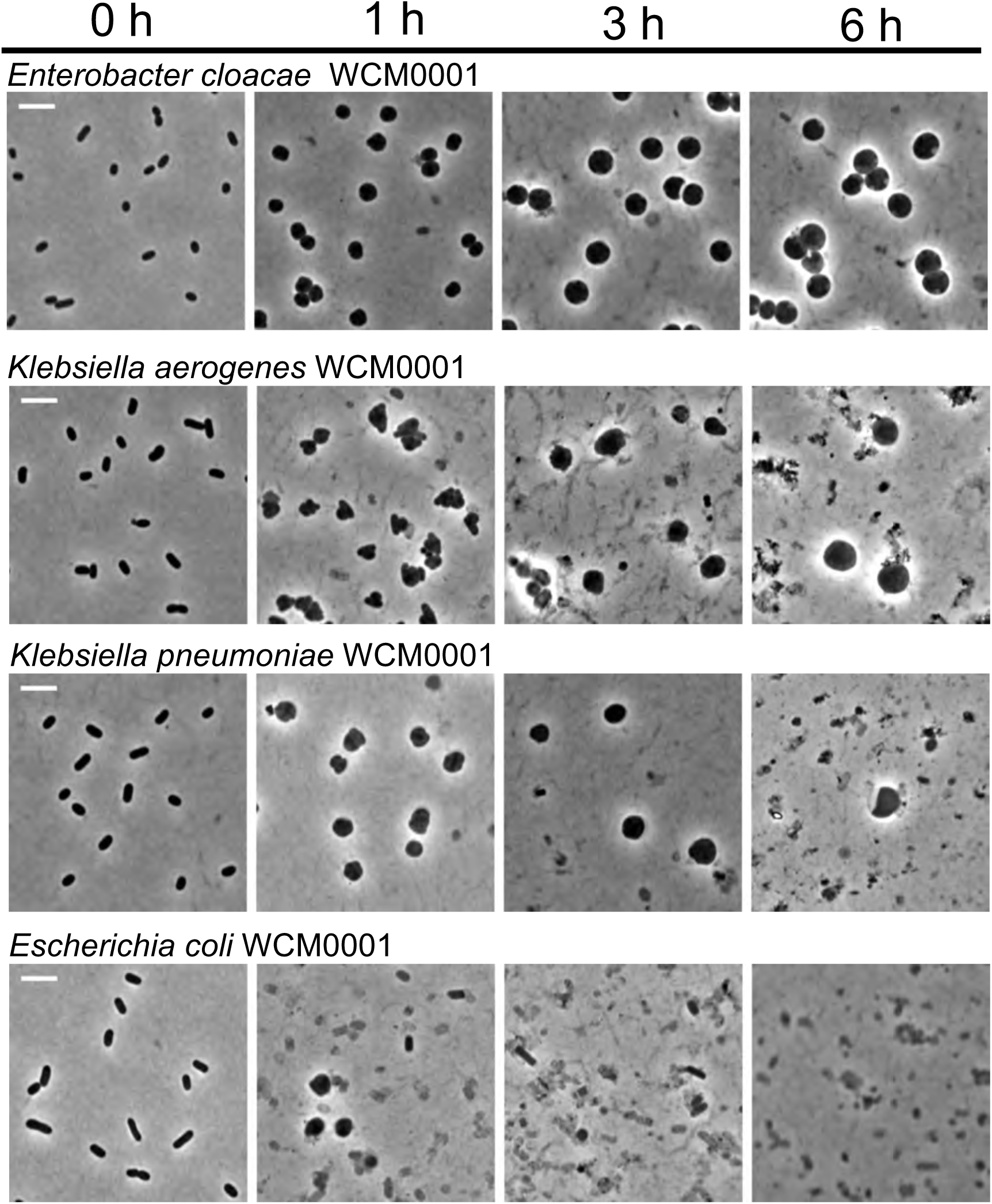
Meropenem exposure induces spheroplast formation in Gram-negative pathogens. Overnight cultures of the indicated isolates were subcultured 1:10 (final volume 5 mL) into pre-warmed BHI+ liquid medium containing 10 µg/mL meropenem and imaged at the indicated time points. Scale bar, 5 µm.

We next used the cell wall stain 7-hydroxycoumarin-amino-D-Alanine (HADA) (30) to test whether the observed spheroplasts were able to survive meropenem exposure by synthesizing cell wall material in a meropenem-insensitive manner, or by being able to sustain structural integrity in the absence of the cell wall. Indeed, osmotically stable, cell wall-containing spherical cells that resemble spheroplasts can be observed in *E. coli* when the elongation-specific class B penicillin-binding protein (PBP) 2 is inhibited (31). Addition of HADA revealed little to no detectable cell wall material in meropenem-treated cells, but, consistent with published data from *E. coli*, did result in strong staining of PBP2-inhibited (*i.e*., mecillinam-treated) cells (Fig. 3). The lack of detectable cell wall material in meropenem-treated cells, combined with their rapid loss of cell shape, suggests these spheroplasts maintain structural integrity through their outer membrane, rather than a reorganized cell wall. This is in line with the recent realization that the Gram-negative OM has a higher than appreciated mechanical load capacity (12). Lastly, loss of the cell wall is typically associated with inhibition of multiple PBPs that include class A PBPs (32). Meropenem has a high affinity for PBP2 (33), but also inhibits multiple other PBPs (PBP1a/b and PBP3) in *E. coli* and *P. aeruginosa* (34, 35) and our data suggest that at the concentration used here, multiple PBPs are inactivated.

**Figure 3.**
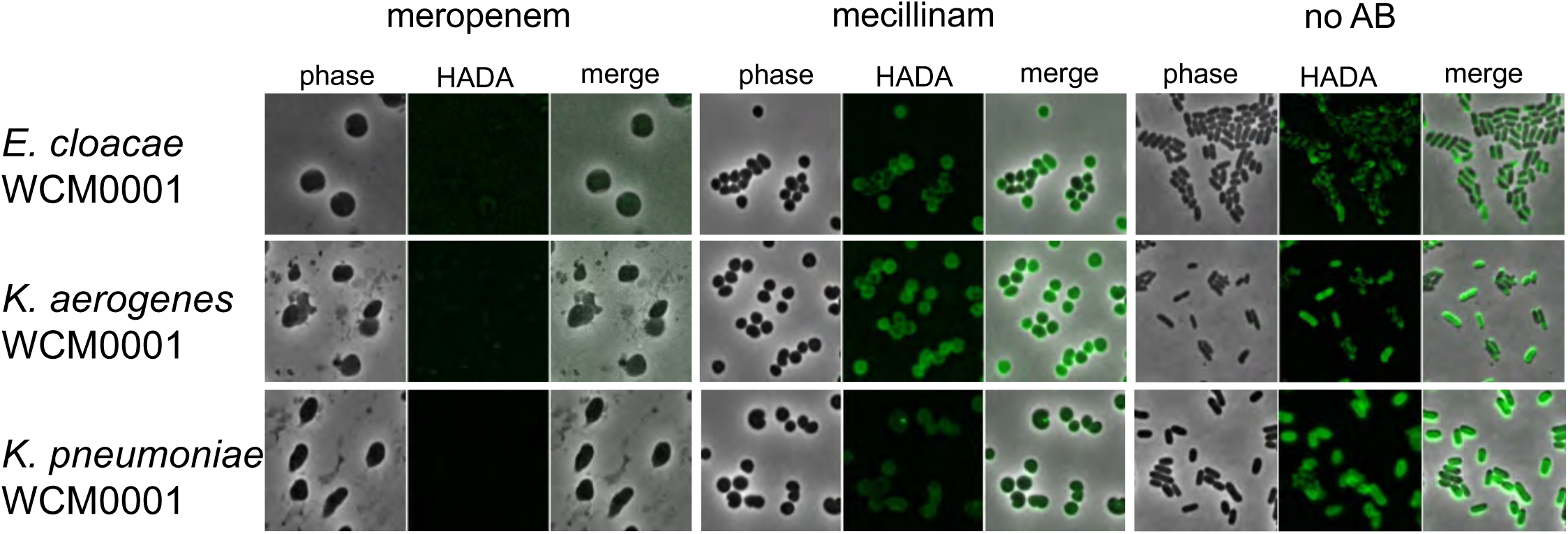
Meropenem-treated spheroplasts have no detectable cell wall material. The indicated isolates were grown in the presence of HADA and treated with either vehicle (no AB), meropenem (10 µg/mL), or mecillinam (20 µg/mL). After 6 hours, cells were washed and imaged using fluorescence microscopy.

If the observed spheroplasts are truly tolerant cells, they should readily revert to rod-shape (*i.e*., wild-type shape) and exponential growth upon removal of the antibiotic. To test this, we withdrew samples after 6 hours of meropenem exposure, removed the antibiotic by addition of purified NDM-1 carbapenemase and imaged these cells in time-lapse. With varying frequencies roughly reflecting the different survival rates, at least some spheroplasts from all isolates were able to recover to rod-shape (Fig. 4, **Fig. S1 – S2, S5-S8**); albeit with different dynamics (cf. *E. cloacae* WCM0001 vs. *K. aerogenes* WCM0001). The recovery process often included rapid division (*e.g*., 25 min after removing the antibiotic in *K. pneumoniae* WCM0001) as spherical cells, resulting in two half-spheroplasts that then increasingly approximated rod-shape during subsequent division events. Taken together, our results suggest that the high tolerance levels observed for the Gram-negative pathogens tested here are not mediated by dormancy, or prolonged lag phase after stationary phase, but rather by the ability to survive for extended time periods without a structurally sound cell wall.

**Figure 4.**
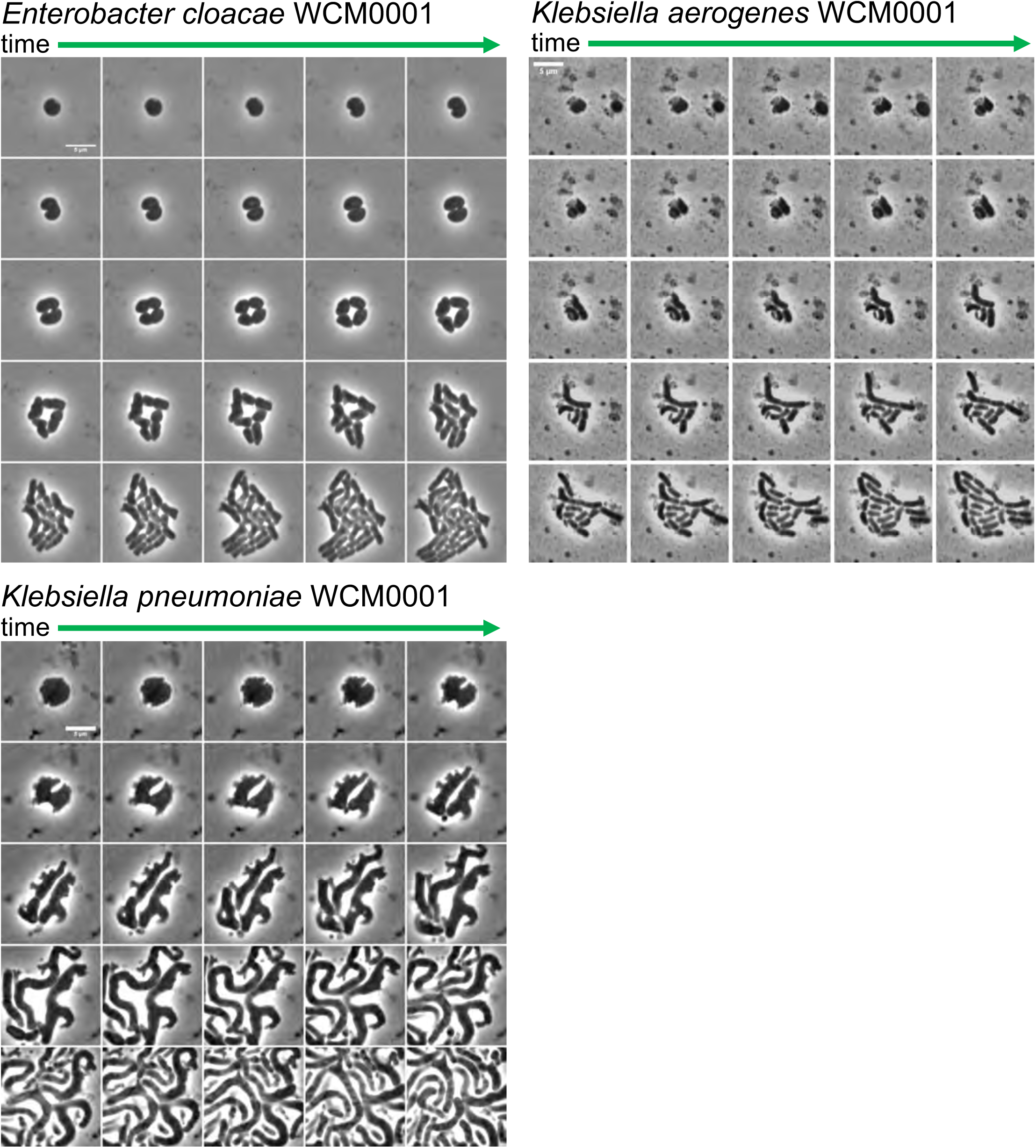
Meropenem-induced spheroplasts can recover to form an exponentially growing population. Time-lapse montage of spheroplasts upon removal of meropenem after 6 hours of treatment. The antibiotic was removed by addition of purified NDM-1 carbapenemase, followed by time-lapse microscopy on BHI+ agarose pads (0.8 % [w/v] agarose). Images were then acquired 5 min apart for another 2 hours. Both *E. coli* isolates were omitted since no spheroplasts were observed after 6 hours of meropenem treatment. Scale bars, 5 µm.

### Quantification of Gram-Negative Tolerance: The Weaver Score

Since our observational data suggested variations in tolerance levels, we sought to quantify the ability to survive and maintain cellular structural integrity during exposure to meropenem. Assays to determine tolerance levels based on killing dynamics have been developed (10, 36); however, we chose to incorporate both cfu/mL and OD_600_ measurements in our tolerance score. In principle, both cfu/mL (viability) and OD_600_ measurements can report on tolerance to β-lactam antibiotics as both are indicators of the bacterial cell’s ability to resist antibiotic-induced lysis. We argue that cell lysis and death can in principle be separable contributors to tolerance. We cannot exclude, for instance, that a significant proportion of antibiotic-damaged spheroplasts die only upon plating on solid media, a condition that results in oxidative stress (37). An isolate that exhibits a significant decrease in viability after exposure to a β-lactam but does not lyse would likely be given a low tolerance designator if cfu/mL only were considered. However, the spheroplasts that do not recover on plates might recover at a high rate under circumstances where cells are not plated following exposure (*e.g*., in the host). Thus, we consider that OD_600_ measurements (in conjunction with cfu/mL) hold informative value for tolerance measurements. We developed a meropenem survival/integrity score, the Weaver tolerance score, by multiplying the fraction of survival (cfu/mL after 6 hours of treatment over initial cfu/mL) with the fraction of OD_600_ readings (OD_600_ reading after 6 hours of treatment over initial OD_600_ reading) (see Methods for details). We were thus able to generate a single value for overall tolerance that considers both parameters of the culture: lysis behavior and plating defect. According to this score, *V. cholerae* N16961*, P. aeruginosa* PA14 and the *E. cloacae* isolates WCM0001 and ATCC 13047 emerged as the most tolerant of the susceptible/non-resistant, non-carbapenemase producing organisms, while both tested *K. pneumoniae* and both *K. aerogenes* isolates exhibited intermediate tolerance, and *E. coli* ranked the lowest (Fig. 5). Consistent with their ability to grow in the presence of meropenem, the KPC-producing isolates scored the highest.

**Figure 5.**
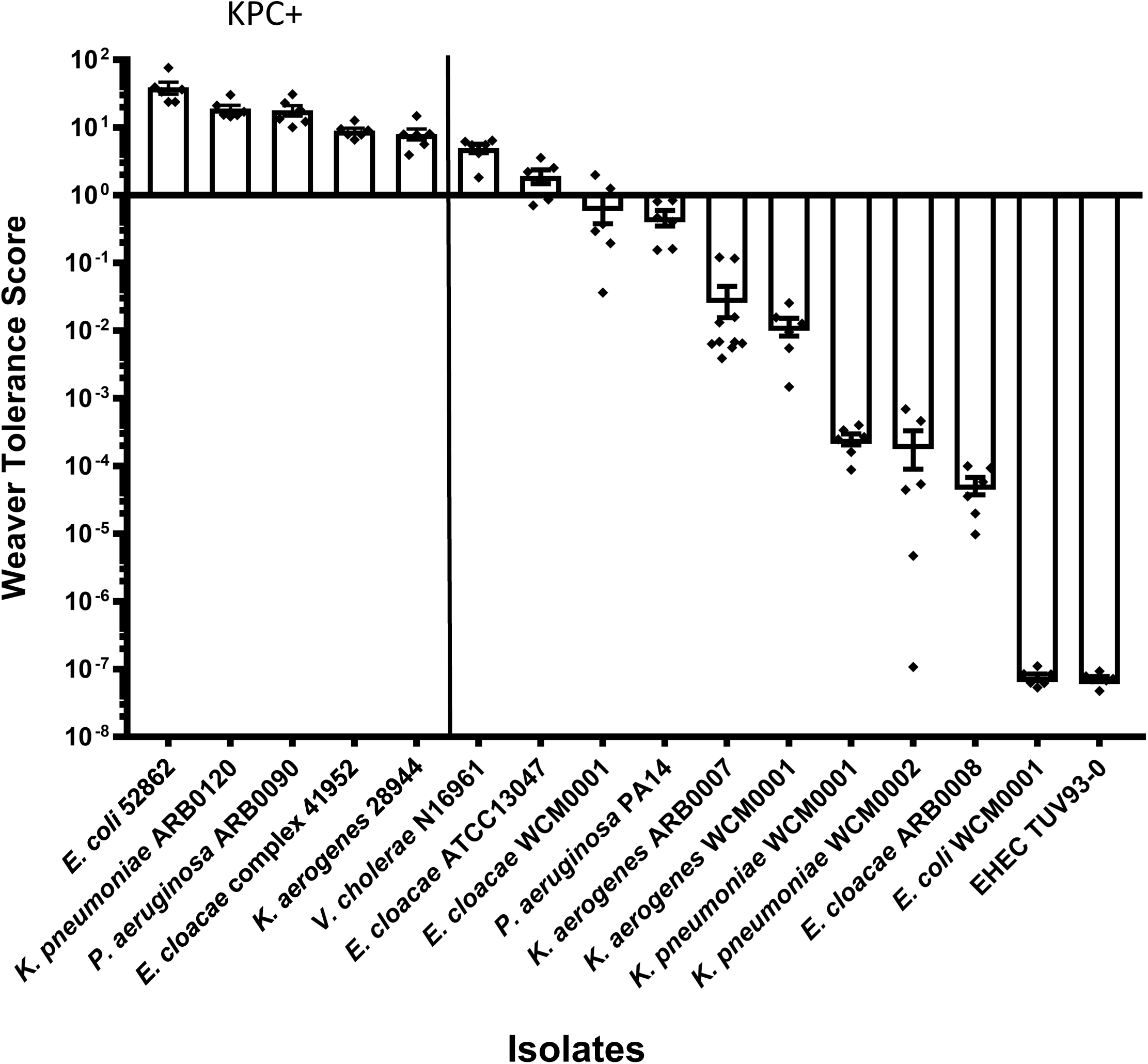
Meropenem Tolerance Scores of Gram-negative Pathogens. Relative tolerance to meropenem was quantified using the Weaver Score equation that is based upon viable cell concentration and optical density data for broth cultures exposed to 10 µg/mL meropenem for 6 hours. The Weaver Score was calculated using the following equation: [(OD_t6_/OD_t0_)*(cfu/mL_t6_/cfu/mL_t0_)], and the resultant values arranged in descending order indicating decreasing tolerance to meropenem. Error bars show standard error of the mean of at least six biological replicates.

### Spheroplast-formation in human serum

To evaluate tolerance in an environment more reminiscent of growth in the human host, we performed killing experiments in human serum. Colony forming units (serum growth medium is incompatible with OD measurements) were measured after six hours of incubation with or without meropenem (Fig. 6) and cells were observed directly for spheroplast-formation. All isolates grew in serum growth medium (Fig. 6B), but compared to BHI+, killing by meropenem was reduced for some isolates. *K. aerogenes* ARB0007 and *K. pneumoniae* WCM0001 were almost completely tolerant with only a ~5-fold decrease in viability over the 6 hour period, compared to the 10 to 100-fold killing that occurs in BHI+ with these isolates. Conversely, *E. cloacae* WCM0001 was killed at a higher rate in human serum than in BHI+. However, spheroplasts were observed for all isolates, except for *E. coli*, and recovery to rod-shape morphology and exponential growth upon removal of the antibiotic by addition of NDM was efficient (Fig. 6C). Taken together, these data suggest that the degree of tolerance is growth-medium specific (though *E. coli* was still the least tolerant), but that spheroplast formation as a means of tolerating exposure to meropenem is conserved across bacteria and conditions tested here.

**Figure 6.**
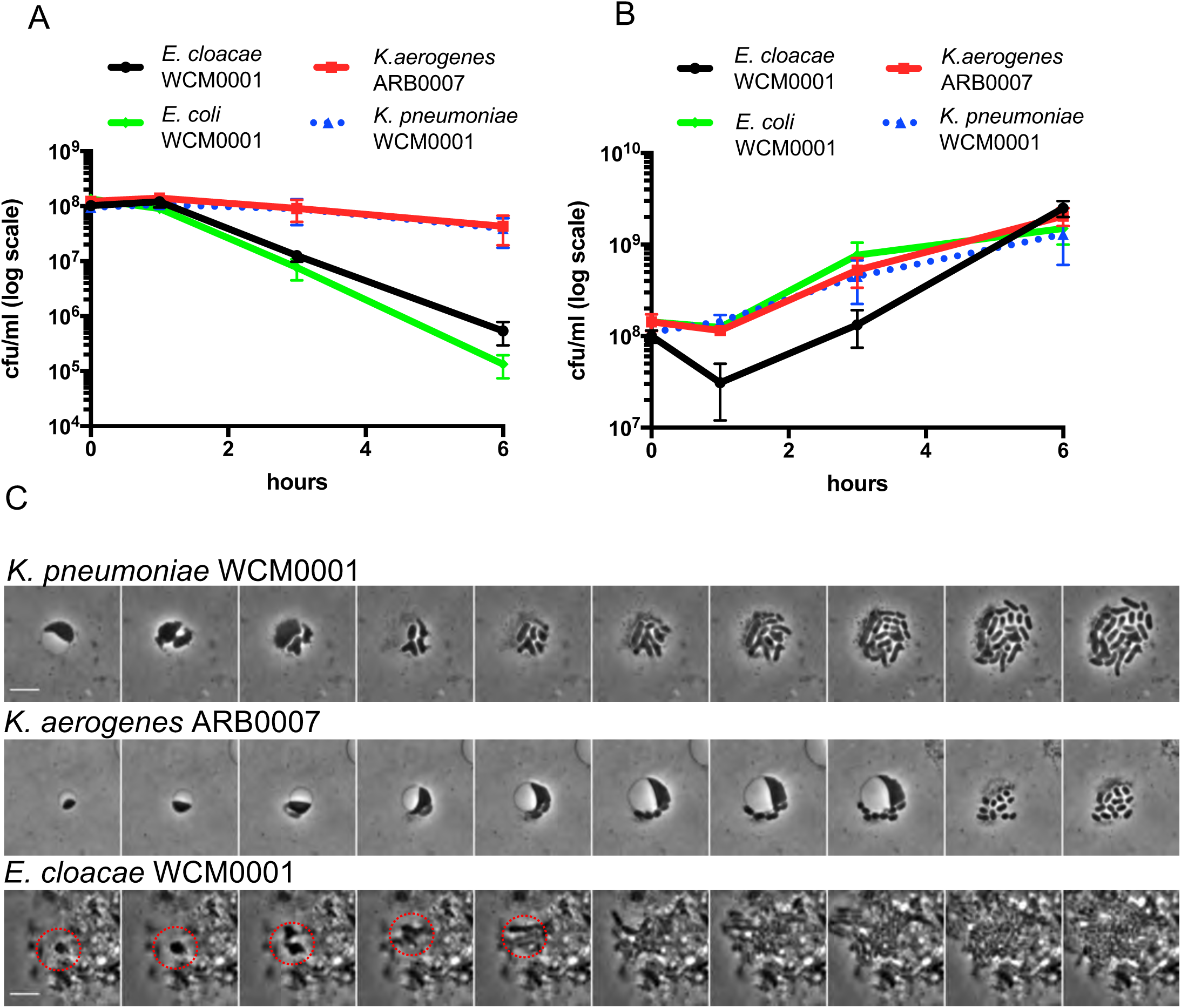
Spheroplast formation and recovery in human serum. The indicated isolates were grown overnight in 40% (v/v) serum/DMEM liquid medium, diluted 10-fold into fresh serum/DMEM liquid medium and incubated in the presence **(A)** or absence **(B)** of 10 µg/mL meropenem. Cells were plated (cfu/mL) at the indicated time points. After 6 hours of incubation, purified NDM-1 carbapenemase was added to remove meropenem, followed by time-lapse microscopy on agarose pads containing 40% (v/v) human serum **(C)**. All values represent the mean of three biological replicates, error bars represent standard error of the mean. Red circle indicates the first steps of a recovering spheroplast within cell debris.

## Discussion

In contrast to antibiotic resistance (the ability to grow in the presence of antibiotics), the phenomenon of antibiotic tolerance (the ability to resist killing by bactericidal antibiotics for extended time periods) remains understudied. While a lot of effort has been directed at understanding persister cells (9) (multidrug tolerant, dormant or near-dormant phenotypic variants produced as a small fraction of bacterial populations), it is unclear what other strategies might exist among bacteria to resist killing by ordinarily bactericidal antibiotics. We and others have previously described a tolerance mechanism (via formation of stable spheroplasts) by which Gram-negative bacteria are able to survive the normally lethal event of cell wall removal that is caused by exposure to β-lactam antibiotics and other inhibitors of cell wall synthesis (13, 14, 16, 17). Cell wall-less Gram-negative spheroplasts are formed by the majority population, are not dormant (*i.e*., not persisters), and presumably rely on their strong OM to maintain structural integrity (16). Here, we show that this phenomenon is more widespread than previously recognized. Gram-negative clinical isolates that are susceptible/non-resistant to meropenem as determined using conventional MIC-based methods failed to be eradicated at appreciable levels of antibiotic, and surviving cells were spheroplasts devoid of detectable cell wall material. Our data raise the possibility that rapid death and lysis of majority populations after β-lactam therapy might be the exception, not the norm, in clinical practice. Therefore in addition to persister formation (8-10, 38), heteroresistance (39), and overt resistance (20), these data highlight a fourth potential mechanism for β-lactam therapy treatment failure.

Importantly, spheroplast-like structures have been isolated from patients that were infected with Gram-negative pathogens and treated with β-lactam antibiotics (3), raising the possibility that these cells are able to survive without a cell wall in the human host. Consistent with this idea, we have observed spheroplast formation in our collection of *Enterobacteriaceae* during exposure to meropenem in human serum growth medium. Therefore, we consider it likely that spheroplasts, similar to persisters, can be responsible for recalcitrant infections. Indeed, herein, we present the first evidence that carbapenem-induced spheroplasts in significant pathogens of the family *Enterobacteriaceae* can readily revert to rod-shape and exponential growth upon removal of carbapenem. While in some previous studies spheroplast formation has been noted (3,13)(40, 41), evidence of spheroplast recovery (*i.e*., reversion to rod-shaped, exponentially growing cells) was lacking. As such, spheroplast formation has not been incorporated into models of antibiotic susceptibility, particularly in the host. In addition, Gram-negative bacteria typically used as model organisms in the laboratory lyse upon exposure to β-lactams in standard laboratory media (*e.g*., LB broth) (42), creating the potential misconception that the standard response to inhibition of cell wall synthesis is lysis. Future experiments will determine if spheroplasts that are able to revert to a growing population are also observed during infections in patients treated with β-lactam antibiotics, and if their formation correlates with treatment outcomes.

In addition to providing a reservoir for a large number of cells that can repopulate an infection after antibiotic therapy is discontinued, a majority population of damaged but recoverable cells, such as spheroplasts after antibiotic treatment, poses other health risks. β-lactam antibiotics have been suggested to induce the generation of reactive oxygen species as well as the SOS DNA damage response, and could thus have mutagenic potential (14,15). A large reservoir of damaged cells might therefore enhance the possibility of developing broad resistance to other antibiotics (16). Furthermore, though we did not test this directly, spheroplasts could in principle continue to produce virulence factors (toxins, proteases and other tissue damaging enzymes), and thus disease, during antibiotic therapy.

The spheroplasts observed here are reminiscent of L-forms in Gram-positive bacteria. However, unlike L-forms, Gram-negative spheroplasts fail to proliferate in the presence of antibiotics and only recover to rod-shape and exponential growth when the antibiotic is removed. L-form cell division relies on membrane lipid overproduction and subsequent stochastic blebbing (43). We speculate that in Gram-negative spheroplasts, L-form-like proliferation is prevented due to their cytoplasmic membranes being confined by their rigid OMs, thus preventing division through membrane blebs.

In summary, our work demonstrates that the ability to survive bactericidal β-lactam antibiotics does not solely rely on classical resistance or dormancy, but instead could also be dependent upon an intrinsic tolerance mechanism in Gram-negative pathogens that are otherwise fully susceptible to the damage induced by cell wall acting agents. Rather than preventing harmful effects of antibiotics (as in resistance and dormancy), these tolerant spheroplasts survive by circumventing the essentiality of the antibiotic’s main target, the cell wall. Our observations underscore the necessity of studying clinical isolates to gain a more complete understanding of the complex processes underlying the susceptibility to antibiotics in clinical settings.

## Acknowledgements

Research in the Dörr lab is supported by NIAID grant 1R01AI143704. Research in the VanNieuwenhze lab is supported by NIH grant GM113172. Isolates *E. cloacae* ARB0008, *K. aerogenes* ARB0007, *K. pneumoniae* ARB0120 and *P. aeruginosa* ARB0090 were obtained from the CDC and FDA Antibiotic Resistance Isolate Bank (https://www.cdc.gov/drugresistance/resistance-bank/index.html). Isolates *E. cloacae* complex 41952, *E. coli* 52862, and *K. aerogenes* 28944 were a gift from Barry Kreiswirth (Rutgers University, NJ). We thank Matthew K. Waldor (Harvard Medical School, MA) and Pamela V. Chang (Cornell University, NY) for the gift of *V. cholerae* N16961 and *E. coli* TUV93-0, respectively.

## Material and Methods

### Chemicals, media and growth conditions

Meropenem (TCI chemicals, Portland, OR) was formulated as a 10 mg/mL stock solution in distilled water and stored at −20°C. BHI+ medium (per liter: 17.5 g brain heart infusion from solids, 10.0 g pancreatic digest of gelatin, 2.0 g dextrose, 5.0 g sodium chloride, 2.5 g disodium phosphate, 15.0 μg/mL hemin, 15.0 μg/mL NAD+) was purchased from RPI (Wilmington, NC) and prepared as a broth according to package instructions; with 15 g per liter agar added for solid media. All isolates were grown in supplemented BHI (BHI+) by adding nicotinamide adenine dinucleotide (NAD; Sigma-Aldrich, St. Louis, MO) and hemin (Beantown Chemical, Hudson, NH), both at a final concentration of 15 µg/mL. All isolates were grown overnight in a 37°C shaking incubator prior to initiating the experiment.

### Bacterial isolates

Bacterial isolates are summarized in Table 1. Clinical isolates were identified to the species or complex level (in the case of *E. cloacae*) using matrix-assisted laser desorption/ionization-time of flight mass spectrometry (MALDI Biotyper, Bruker Daltonics, Inc., Billerica, MA) according to the manufacturer’s instructions. Meropenem AST (Table 1) was performed for all isolates using gradient diffusion (Etest, bioMérieux, Inc., Durham, NC) according to the manufacturer’s instructions on both MHA and BHI+ agar (*i.e*., solid media). Each day of testing, quality control testing of the Etest strips was performed on both MHA and BHI+ with *E. coli* ATCC 25922. In all cases, quality control passed on both MHA and BHI+ agar. The resultant AST data obtained with MHA was interpreted using the CLSI M100-S29 (*Enterobacteriaceae* and *P. aeruginosa*) and M45 (*V. cholerae*) documents (24, 25). For the KPC-producing isolates the presence of the *bla*KPC gene was confirmed using the Xpert Carba-R assay (Cepheid, Sunnyvale, CA, USA) according to the manufacturer’s instructions.

### Microscopy

All images were taken on a Leica MDi8 microscope (Leica Microsystems, GmbH, Wetzlar, Germany) with a PECON TempController 2000-1 (Erbach, Germany), heated stage at 37 °C for growth experiments, or room temperature for static images. Time-lapse microscopy was performed by imaging frames five minutes apart and data were processed in ImageJ (44). HADA stained cells were imaged at 365 nm excitation for one second exposure time. Images were minimally processed in ImageJ by subtracting background and adjusting brightness/contrast uniformly across all fluorescent images.

### Time-dependent killing experiments

Overnight cultures of each isolate were grown at 37°C in liquid BHI+ medium and the following day diluted 1:10 in fresh, pre-warmed BHI+ medium containing a final concentration of 10 µg/mL meropenem. At each time-point, samples were diluted five-fold in blank medium and OD_600_ was measured. At the same time point, viable cell counts were also assessed by 10-fold serially diluting cells in BHI+ and spot-plating 10 µL of each dilution on BHI+ agar plates. Colonies were counted after 24 hours growth at 30°C. Images were taken by placing cells on BHI+ agarose pads (0.8 % [w/v] agarose). Cells were concentrated by centrifugation (8,000 *x g*, 5 min) where necessary.

### Weaver tolerance score

The tolerance score was calculated from measurements of OD_600_ and cfu/mL after 6 hours of exposure to meropenem. The score was calculated as (OD_t6_/OD_t0_) × (cfu/mL_t6_/cfu/mL_t0_); *i.e*., OD-fold change multiplied with survival fraction.

### Time-dependent killing assays in human serum

To generate serum growth medium (SGM), human serum (Rockland Pharmaceuticals, Limerick, PA) was thawed on ice and diluted in Dulbecco’s modified Eagle’s medium (DMEM, VWR, Radnor, PA) to 40% (v/v). Bacteria were inoculated from frozen stocks into 300 µL of SGM in Eppendorf tubes and incubated at 37 °C overnight without agitation. After incubation, cells were diluted 10-fold into 450 µL of fresh SGM, followed by addition of meropenem (10 µg/mL). Survival was measured by diluting and spot plating for cfu/mL at the indicated times. For recovery time-lapse images, cells were concentrated 10-fold (via centrifugation, 8,000 rcf for 5 min) and the antibiotic inactivated by addition of 5 µL of purified NDM-1 (5 mg/mL). Time-lapse images were obtained at 37°C on SGM + 0.8% (w/v) agarose.

### HADA staining following antibiotic treatment

Cultures were grown shaking at 37°C BHI+ liquid media and subcultured the next day 1:10 to total 1 mL volumes containing 50 µM HADA (30) (7-hydroxycoumarin-amino-D-Alanine) with or without meropenem (10 µg/mL). At each time-point, 100 µL of the culture was harvested and washed three times with 200 µL BHI+ by centrifugation (5 min, 8000 *x g*) to remove antibiotic and excess HADA. After the third wash, cells were concentrated 10-fold and imaged on BHI+ agarose pads (0.8 % [w/v] agarose). Where indicated, HADA staining/imaging was performed as described above after treatment with 20 µg/mL mecillinam (Sigma-Aldrich, St. Louis, MO). Images were analyzed in ImageJ and are minimally processed (background removal).

### Purification of New Delhi Metallo-β-lactamase-1 (NDM-1)

Isolate *E. cloacae* ATCC BAA-2468 was used as a template for the PCR-amplification of the *blaNDM-1* gene. SignalP 4.1 was used to predict the membrane-localization signal sequence of *NDM-1*. PCR primers BR_83 (5′-cagcagcggcctggtgccgcgcggcagccaGTGCATGCCCGGTGAAATCCG-3′) and BR_84 (5′-cagcttcctttcgggctttgttagcagccgCATGGCTCAGCGCAGCTTGTC-3′) were designed to amplify the gene without the predicted signal sequence. Following PCR using Q5 DNA-polymerase (New England Biolabs, Ipswich, MA) and the BR_83/BR_84 primer pair, the product was cloned into the pET1-5b N-terminal 6×His expression plasmid (New England Biolabs).

The plasmid was transformed by heat-shock into chemically competent *E. coli* BL21 (DE3) cells (New England Biolabs). Using the transformed cells, 1 L LB medium cultures were grown from single colonies shaking at 37°C. At OD_600_ ~ 0.3, cells were induced with 1 mM isopropyl-β-D-thiogalactopyranoside (Sigma-Aldrich, St. Louis, MO) and grown for an additional 3 hours at 37°C. Cells were harvested by centrifugation (20 min, 11,200 *xg*) and the pellets frozen at −80°C. After lysis by sonication, the protein was found to be insoluble. Insoluble protein pellets were resolubilized in 3 M urea (VWR, Radnor, PA) and purified using immobilized metal affinity chromatography using Ni-NTA resin (Qiagen, Hilden, Germany). Eluted proteins were renatured by three-step dialysis to a final buffer composition of 20 mM Tris, 150 mM NaCl, 50 µM ZnSO_4_ and 30% (v/v) glycerol. The resulting protein was quantified by Bradford Assay (45) and its functionality verified in a biological assay by purified NDM-1’s ability to restore growth of meropenem-susceptible *E. coli* MG1655 on agar containing meropenem (10 µg/mL).

## Supplemental Figure Legends

**Figure S1. Spheroplast formation and recovery in *V. cholerae* N16961**

**(A)** Survival (cfu/mL, blue; OD_600_, red) in the presence of 10 µg/mL meropenem; error bars show the standard error of the mean of three biological replicates. **(B)** Spheroplast formation at 0, 1, 3, and 6 hours after exposure to 10 µg/mL meropenem. **(C)** Time-lapse images showing recovery upon removal of meropenem with purified NDM-1 carbapenemase (frames were acquired 5 min apart). Scale bars, 5 µm.

**Figure S2. Spheroplast formation and recovery in *P. aeruginosa* PA14**

**(A)** Survival (cfu/mL, blue; OD_600_, red) in the presence of 10 µg/mL meropenem, error bars show the standard error of the mean of three biological replicates. **(B)** Spheroplast formation at 0, 1, 3, and 6 hours after exposure to 10 µg/mL meropenem. **(C)** Time-lapse montage of spheroplasts upon removal of meropenem after 6 hours of treatment. The antibiotic was removed by addition of purified NDM-1 carbapenemase, followed by time-lapse microscopy on BHI+ agarose pads (0.8 % [w/v] agarose). Images were then acquired 5 min apart for another 2 hours. Both *E. coli* isolates were omitted since no spheroplasts were observed after 6 hours of meropenem treatment. Scale bars, 5 µm.

**Figure S3. Growth of susceptible/non-resistant and carbapenemases-producing isolates without antibiotic treatment.**

Growth of the indicated isolates in BHI+ (cfu/mL, solid lines; OD_600_, dotted lines), error bars show standard error of the mean of two biological replicates.

**Figure S4. Growth of KPC-producing *Enterobacteriaceae* in the presence of meropenem.**

Growth of the indicated isolates in BHI+ (cfu/mL, solid lines; OD_600_, dotted lines) supplemented with 10 µg/mL meropenem, error bars show standard error of the mean of two biological replicates.

**Figure S5. Spheroplast formation and recovery in *E. cloacae* ATCC 13047**

**(A)** Spheroplast formation at 0, 1, 3, and 6 hours after exposure to 10 µg/mL meropenem in BHI+. **(B)** Time-lapse montage of spheroplast recovery upon removal of meropenem after 6 hours of treatment. The antibiotic was removed by addition of purified NDM-1 carbapenemase, followed by time-lapse microscopy on BHI+ agarose pads (0.8 % [w/v] agarose). Images were acquired 5 min apart for another 2 hours. Scale bars, 5 µm.

**Figure S6. Spheroplast formation and recovery in *K. aerogenes* ARB0007**

**(A)** Spheroplast formation at 0, 1, 3, and 6 hours after exposure to 10 µg/mL meropenem in BHI+. **(B)** Time-lapse montage of spheroplast recovery upon removal of meropenem after 6 hours of treatment. The antibiotic was removed by addition of purified NDM-1 carbapenemase, followed by time-lapse microscopy on BHI+/0.8% (w/v) agarose. Images were acquired 5 min apart for another 2 hours. Scale bars, 5 µm.

**Figure S7. Spheroplast formation and recovery in *K. pneumoniae* WCM0002**

**(A)** Spheroplast formation at 0, 1, 3, and 6 hours after exposure to 10 µg/mL meropenem. **(B)** Time-lapse montage of spheroplast recovery upon removal of meropenem after 6 hours of treatment. The antibiotic was removed by addition of purified NDM-1 carbapenemase, followed by time-lapse microscopy on BHI+ agarose pads (0.8 % [w/v] agarose). Images were acquired 5 min apart for another 2 hours. Scale bars, 5 µm.

**Figure S8. Spheroplast formation and recovery in *E. cloacae* ARB0008**

**(A)** Spheroplast formation at 0, 1, 3, and 6 hours after exposure to 10 µg/mL meropenem. **(B)** Time-lapse montage of spheroplast recovery upon removal of meropenem after 6 hours of treatment. The antibiotic was removed by addition of purified NDM-1 carbapenemase, followed by time-lapse microscopy on BHI+ agarose pads (0.8 % [w/v] agarose). Images were acquired 5 min apart for another 2 hours. Scale bars, 5 µm.

**Figure S9. Absence of spheroplasts in *E. coli* isolates.**

Overnight cultures of *E. coli* WCM0001 **(A)**, and TUV93-0 **(B)** were subcultured 1:10 (final volume 5 mL) into pre-warmed BHI+ liquid medium supplemented with 10 µg/mL meropenem and imaged at the indicated time points. Scale bar, 5 µm.

**Figure S10. Cell Morphology without antibiotic treatment.**

Overnight cultures of the indicated isolates were diluted 10-fold into BHI+ liquid media, and imaged at the indicated time points. Scale bar, 5 µm.

**Figure S11. Cell Morphology of KPC-producing *Enterobacteriaceae*.**

Overnight cultures of the indicated isolates were diluted 10-fold into fresh BHI+ liquid media containing vehicle **(A)**, or meropenem (10 µg/mL) **(B)**, and imaged at the indicated time points. Scale bar, 5 µm.

