## Supplemental Figures for "Spheroplast-mediated carbapenem tolerance in Gram-negative pathogens"

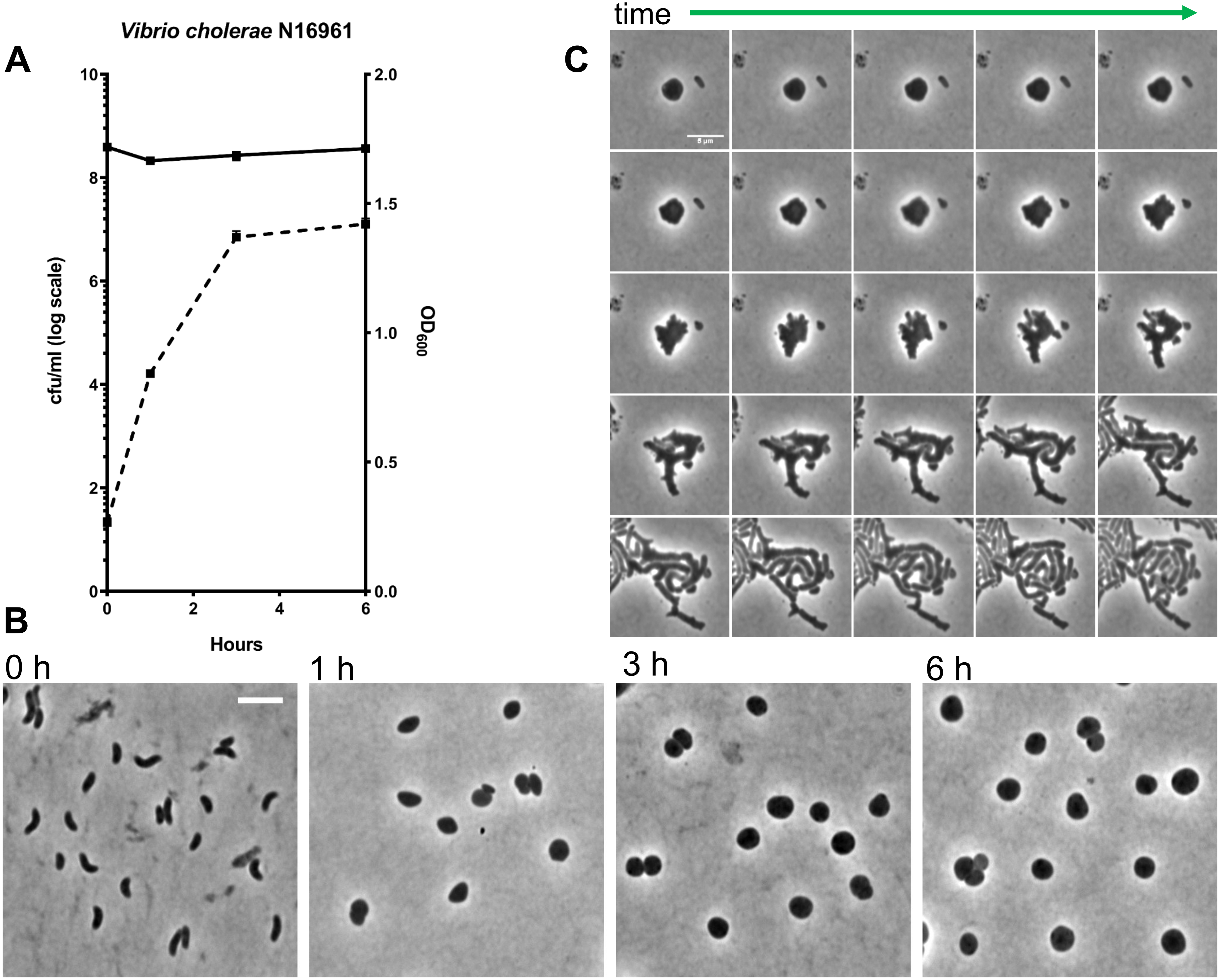
A
Vibrio cholerae N16961
cfu/ml (log scale)
OD600
Hours
0
2
4
6
8
10
0.0
0.5
1.0
1.5
2.0
B
0 h
1 h
3 h
6 h
C
time

**Fig. S1**

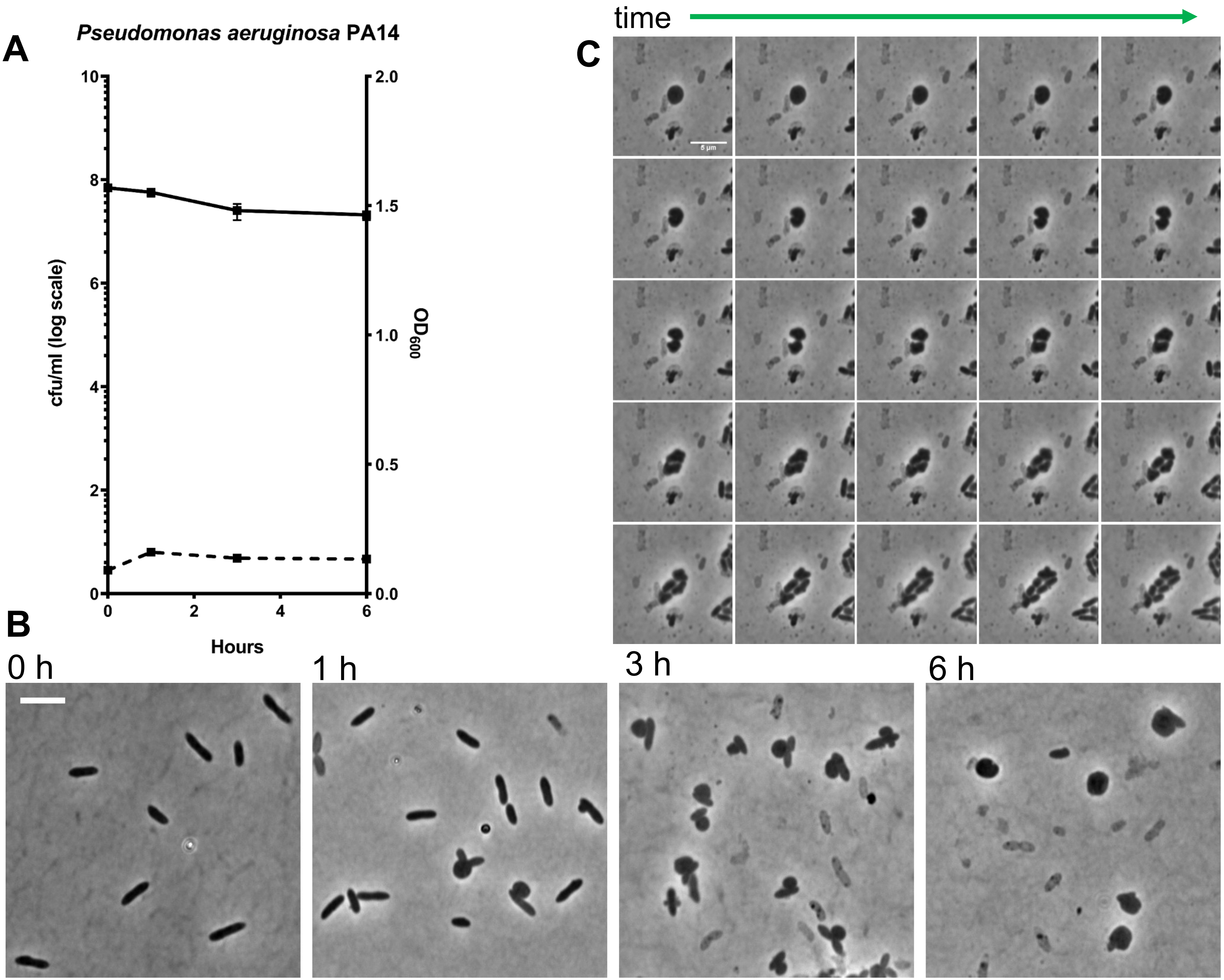
A
Pseudomonas aeruginosa PA14
cfu/ml (log scale)
OD600
Hours
0
2
4
6
8
10
0.0
0.5
1.0
1.5
2.0
B
0 h
1 h
3 h
6 h
C
time
5 µm

Fig. S2

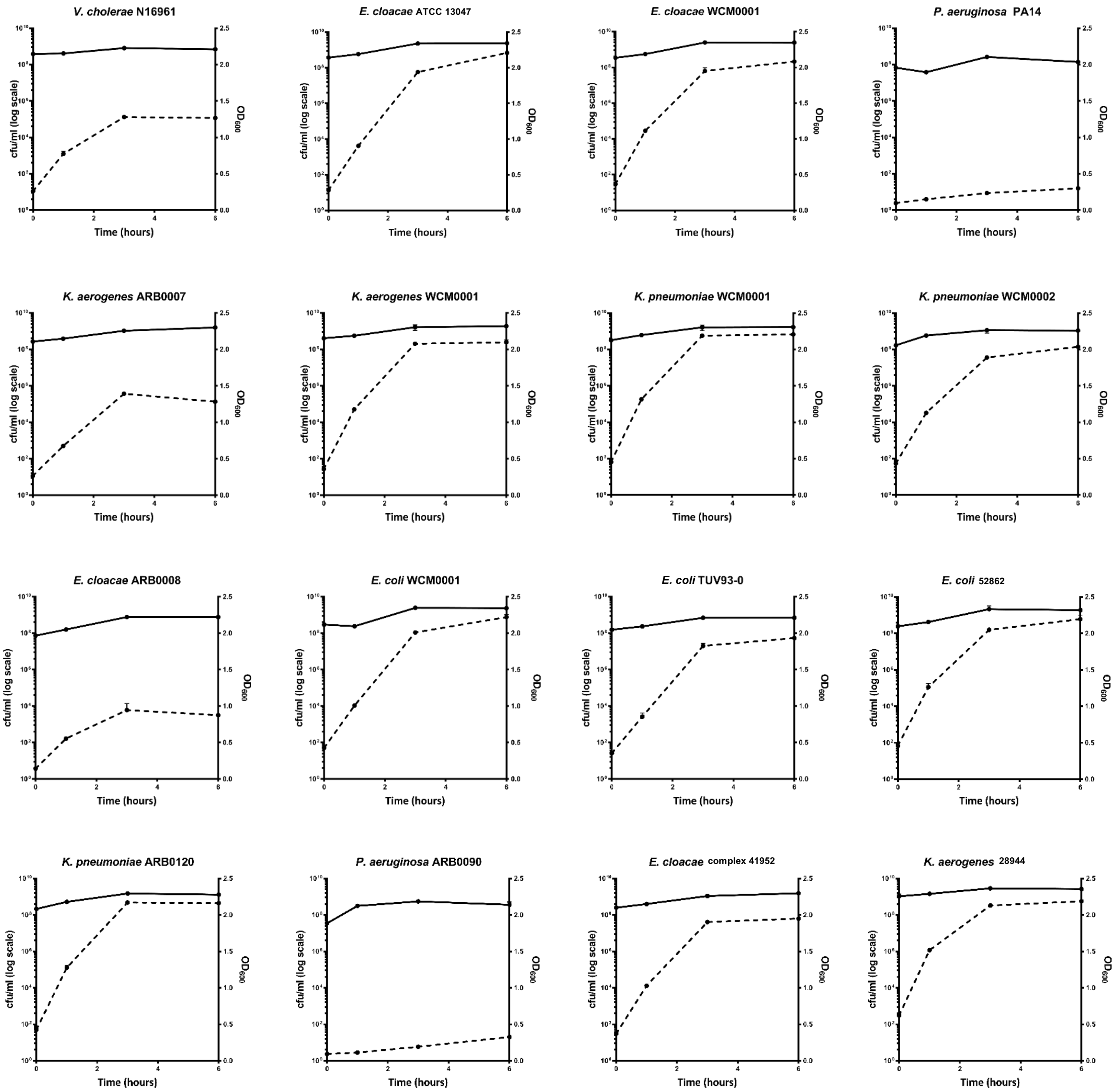
V. cholerae N16961
E. cloacae ATCC 13047
E. cloacae WCM0001
P. aeruginosa PA14
K. aerogenes ARB0007
K. aerogenes WCM0001
K. pneumoniae WCM0001
K. pneumoniae WCM0002
E. cloacae ARB0008
E. coli WCM0001
E. coli TUV93-0
E. coli 52862
K. pneumoniae ARB0120
P. aeruginosa ARB0090
E. cloacae complex 41952
K. aerogenes 28944
cfu/ml (log scale)
$OD_{600}$
Time (hours)

**Fig. S3**

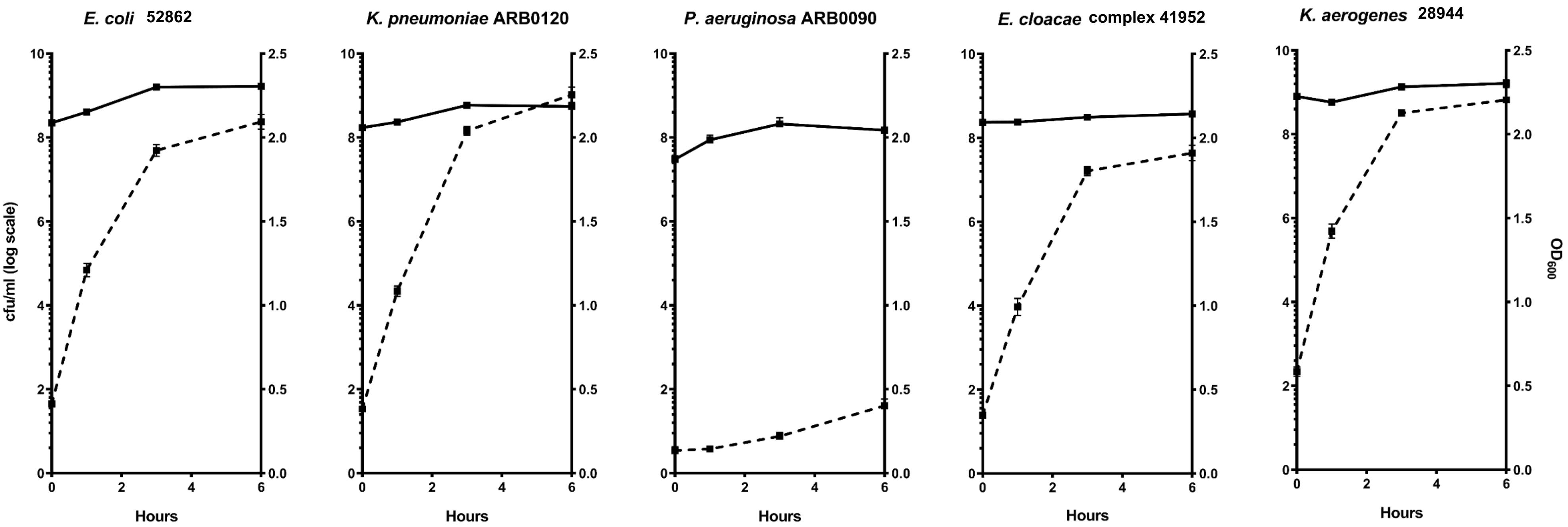

E. coli 52862
K. pneumoniae ARB0120
P. aeruginosa ARB0090
E. cloacae complex 41952
K. aerogenes 28944
cfu/ml (log scale)
OD600
Hours


### Fig. S4

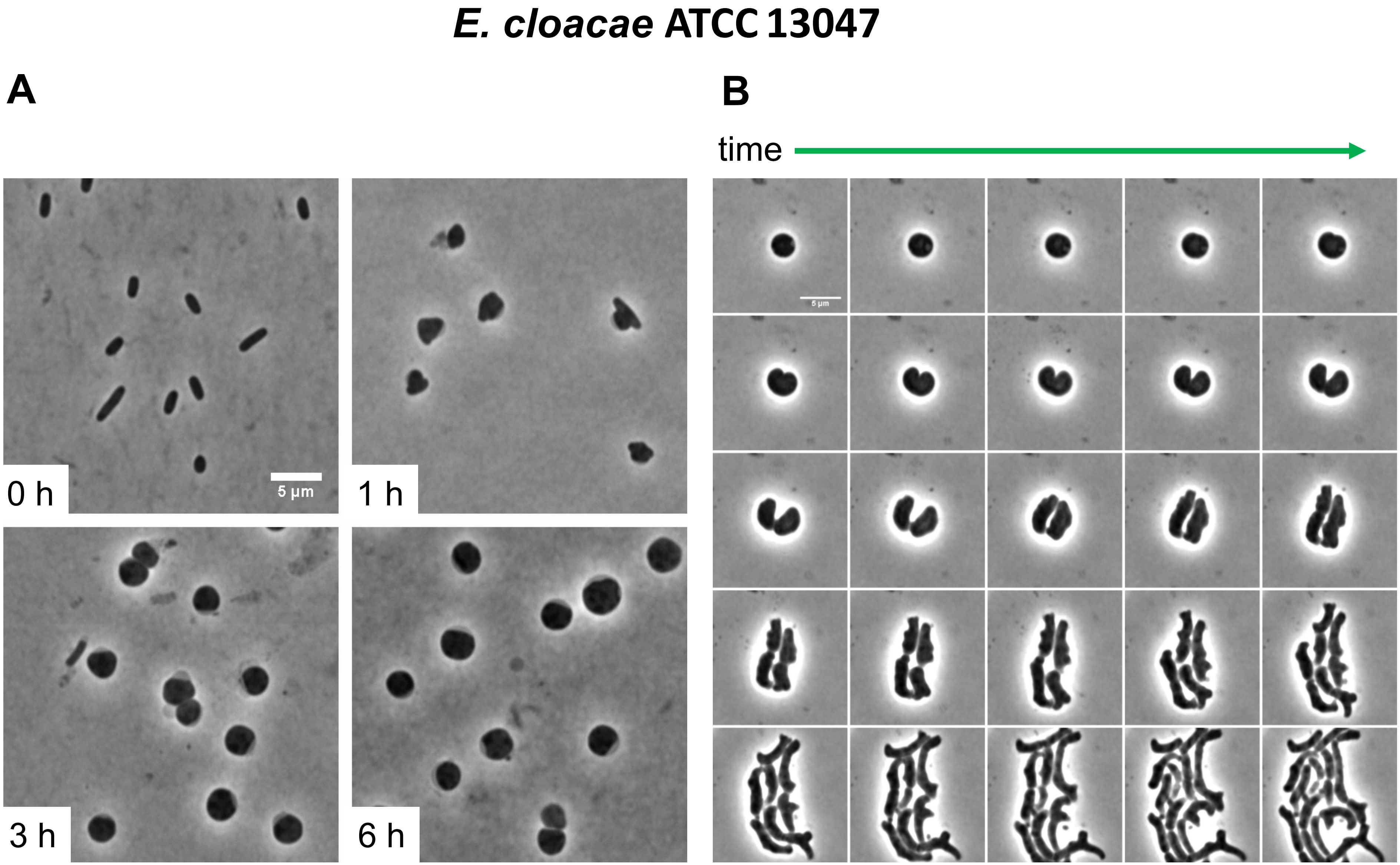


**Fig. S5**

#### *K. aerogenes* ARB0007

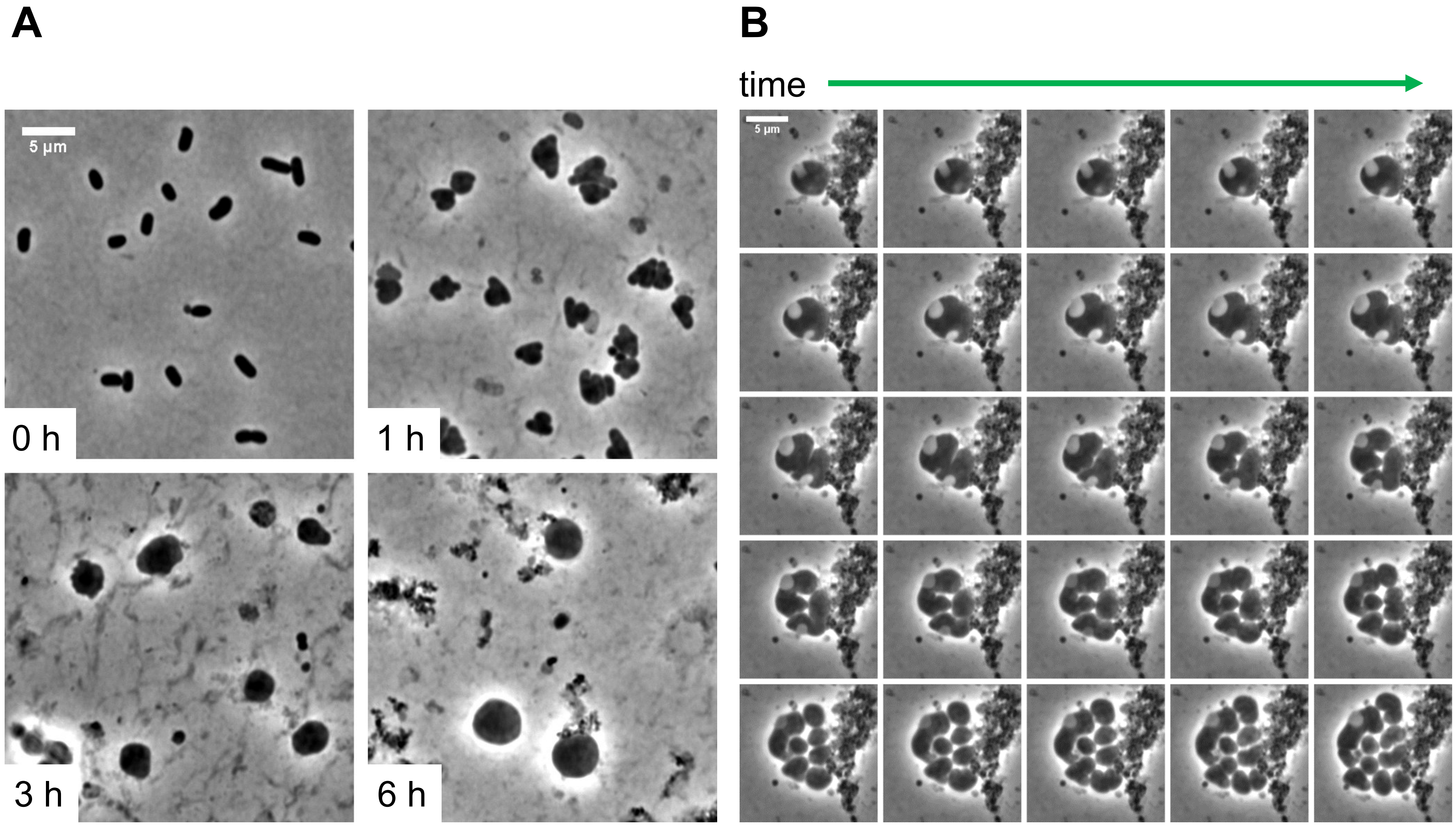


**Fig. S6**

### *K. pneumoniae* WCM0002

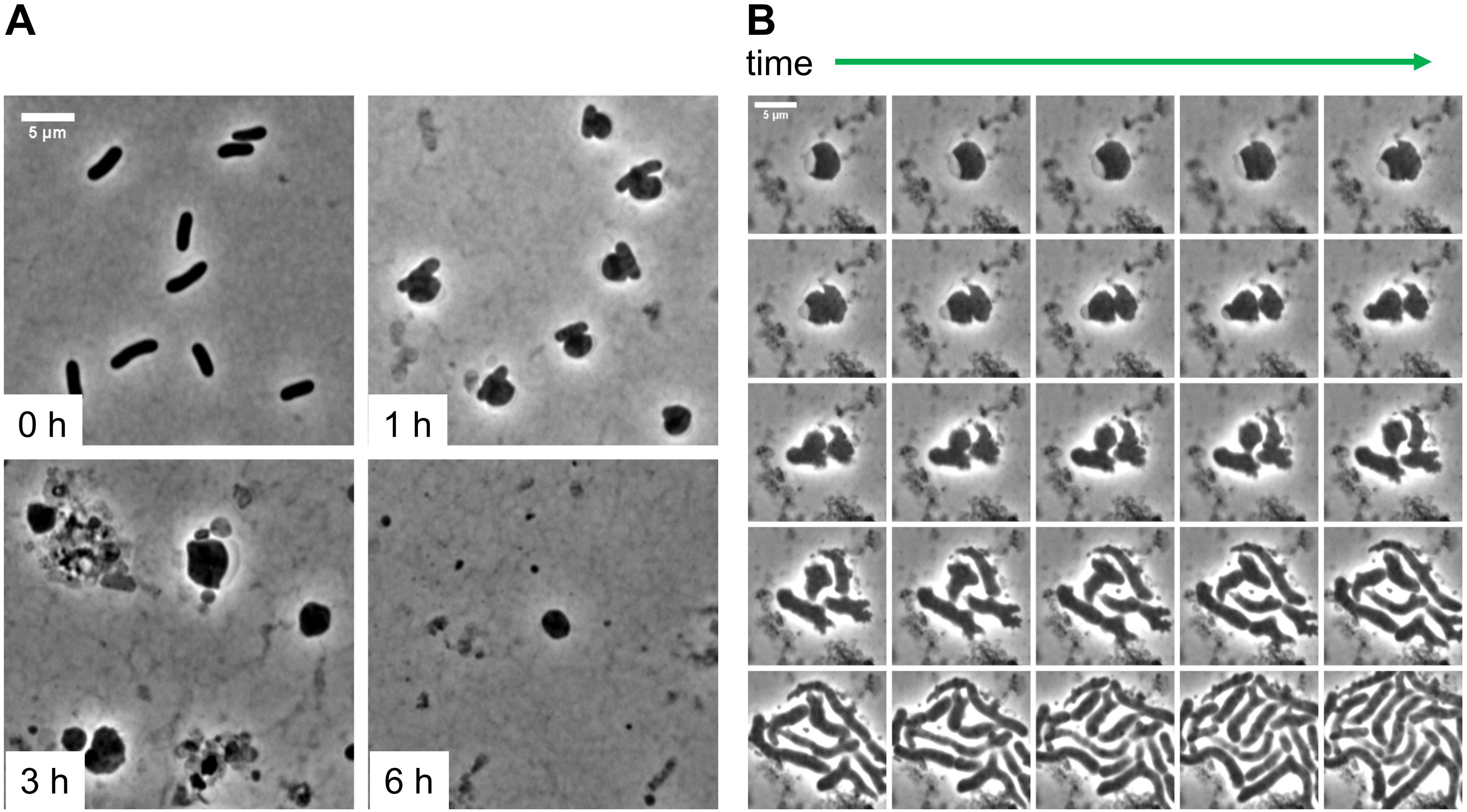


**Fig. S7**

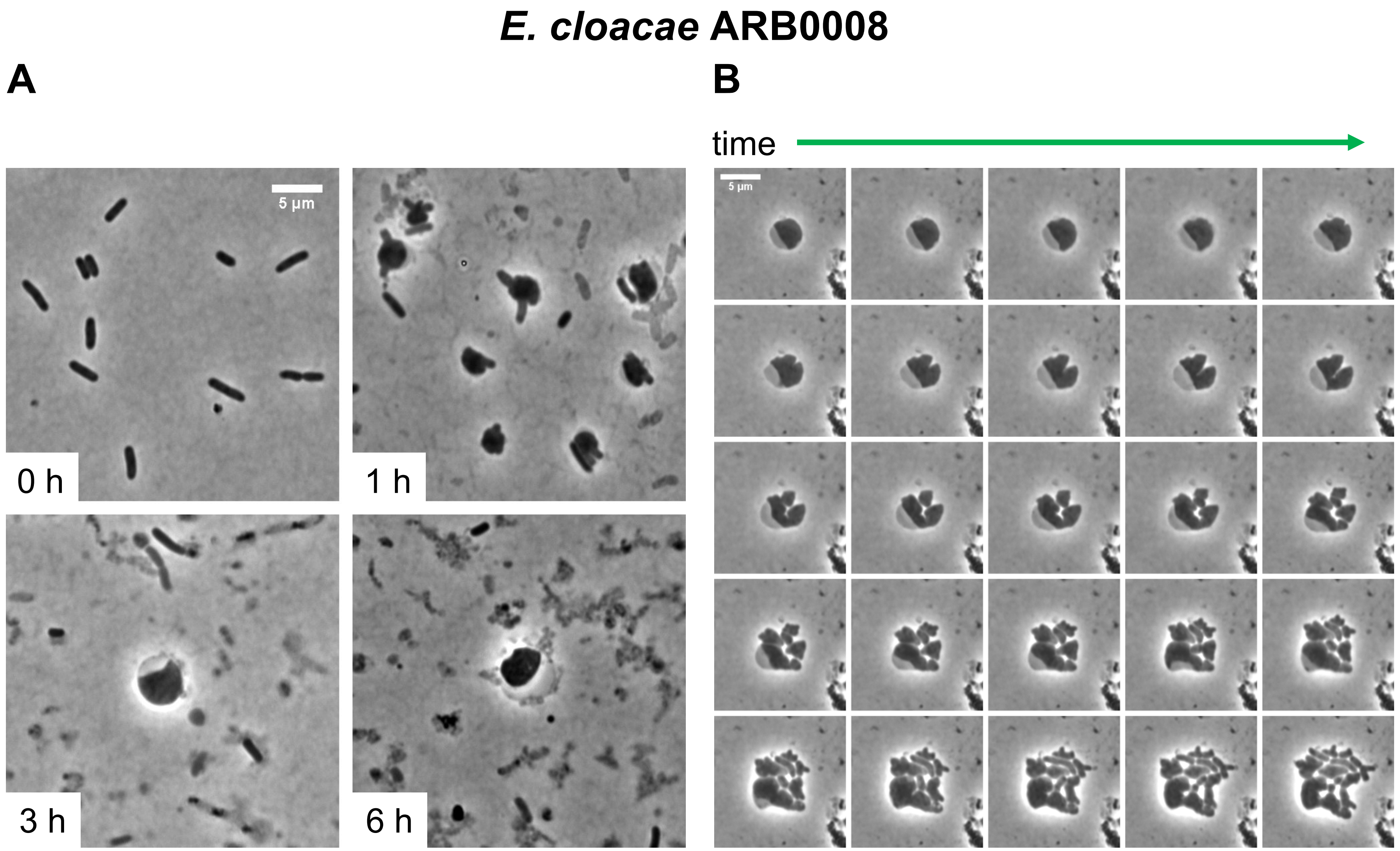


Fig. S8

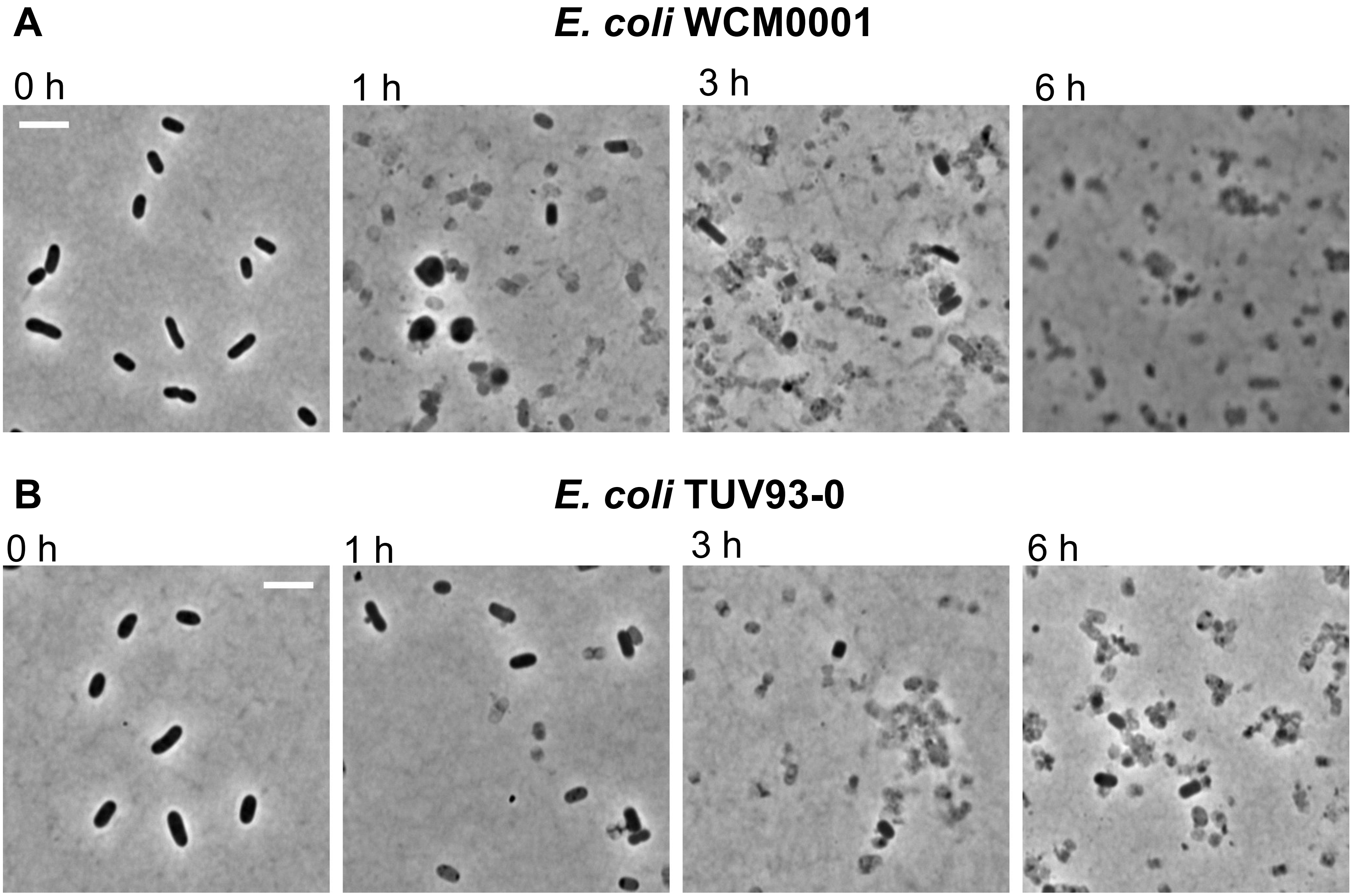


Fig. S9

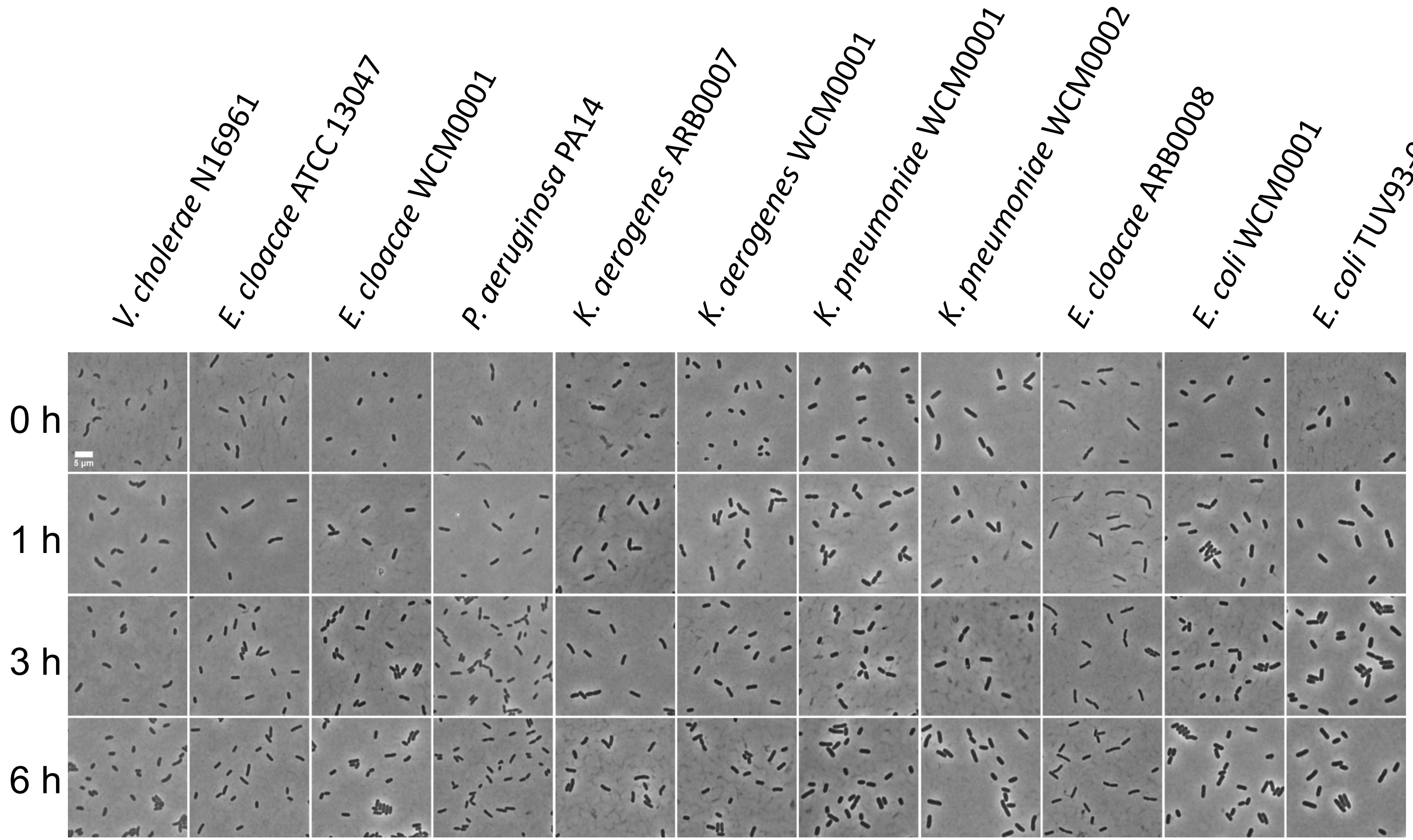
V. cholerae N16961
E. cloacae ATCC 13047
E. cloacae WCM0001
P. aeruginosa PA14
K. aerogenes ARB0007
K. aerogenes WCM0001
K. pneumoniae WCM0001
K. pneumoniae WCM0002
E. cloacae ARB0008
E. coli WCM0001
E. coli TUV93-0
0 h
1 h
3 h
6 h
5 μm

**Fig. S10**

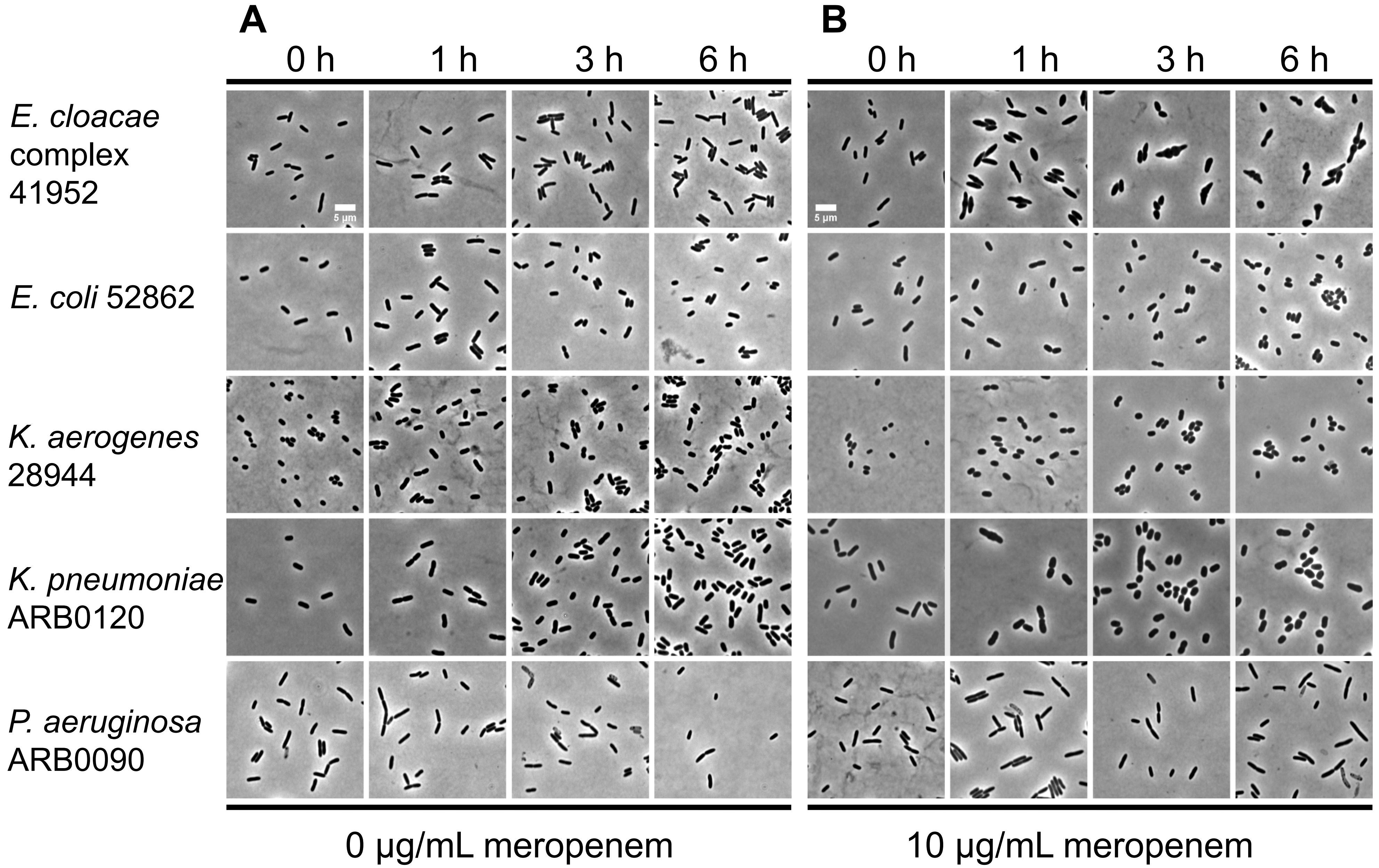
A
B
0 h
1 h
3 h
6 h
0 h
1 h
3 h
6 h
E. cloacae complex 41952
E. coli 52862
K. aerogenes 28944
K. pneumoniae ARB0120
P. aeruginosa ARB0090
5 µm
5 µm
0 µg/mL meropenem
10 µg/mL meropenem

**Fig. S11**
